# Targeting cellular cathepsins inhibits hepatitis E virus infection

**DOI:** 10.1101/2023.11.03.565430

**Authors:** Mara Klöhn, Thomas Burkard, Juliana Janzen, Jil Alexandra Schrader, André Gömer, Richard J. P. Brown, Viet Loan Dao Thi, Volker Kinast, Yannick Brüggemann, Daniel Todt, Eike Steinmann

**Author notes:** equal contribution (Mara Klöhn, Thomas Burkard). Corresponding author: Prof. Dr. Eike Steinmann Department of Molecular and Medical Virology, Ruhr-University Bochum, Bochum, Germany Universitätsstr. 150, 44801 Bochum.

## Abstract

**Background and Aims:** The hepatitis E virus (HEV) is estimated to be responsible for 70,000 deaths annually, yet therapy options remain limited. In the pursuit of effective antiviral therapies, targeting viral entry holds promise and has proven effective for other hepatotropic viruses. However, the precise mechanisms and host factors required during HEV entry remain unclear. Cellular proteases have emerged as a class of host factors required for viral surface protein activation and productive cell entry by many viruses. Hence, we investigated the functional requirement and therapeutic potentials of cellular proteases during HEV infection.

**Approach and Results:** Using our recently established HEV cell culture model and subgenomic HEV replicons, we found that blocking lysosomal cathepsins (CTS) with small molecule inhibitors, impedes HEV infection without affecting replication. Most importantly, the pan-cathepsin inhibitor K11777 robustly suppressed HEV infections with an EC_50_ of ∼ 0.01 nM. Inhibition by K11777, devoid of notable toxicity in hepatoma cells until micromolar concentrations, was also observed in differentiated HepaRG and *ex vivo* in primary human hepatocytes. Furthermore, through time-of-addition experiments, we confirmed that HEV entry is potently blocked by inhibition of cathepsins and cathepsin L (CTSL) knockout cells were less permissive to HEV suggesting that CTSL is critical for HEV infection.

**Conclusions:** In summary, our study highlights the pivotal role of lysosomal cathepsins, especially CTSL, in the HEV entry process. The profound anti-HEV efficacy of the pan-cathepsin inhibitor, K11777, especially with its notable safety profile in primary cells, further underscores its potential as a promising therapeutic candidate.

## Introduction

Hepatitis E virus (HEV) is the main cause of acute hepatitis worldwide, affecting at least 20 million people annually, leading to 3.3 million symptomatic infections and approximately 70,000 fatalities (1). While HEV infections often remain asymptomatic, they can progress to chronic disease in immunocompromised patients, notably transplant recipients and cancer patients, resulting in rapid progression to cirrhosis and liver failure (2). Current therapeutic options against HEV are limited to the off-label use of the broad-spectrum antiviral agent ribavirin (RBV) and pegylated Interferon-alpha (pegIFNα) (3). However, RBV therapy is often discontinued due to adverse side effects and is only effective in ∼80% of patients, while 20% of treated patients remain viremic (3,4).

HEV is a member of the *Hepeviridae* family and has a 7.2 kB single-stranded RNA genome with positive polarity (5,6). The genome of HEV encompasses three open reading frames (ORFs) which encode for: a nonstructural polyprotein (ORF1) essential for viral replication (7), the capsid protein (ORF2) which undergoes endoplasmic recycling (8–10) and a small ORF3 protein with multi-regulatory functions (11–13). HEV is a quasi-enveloped virus; it exists either as a non-enveloped form (neHEV) in bile or circulates as a host-cell lipid-derived enveloped form (eHEV) in the blood (14). Historically, efficient *in vitro* culture systems have been lacking, leaving many facets of the HEV life cycle including virus-host interactions that facilitate attachment to cells and/or entry largely elusive (15). However, current evidence suggests distinct entry mechanisms for both HEV forms (16). Notably, studies using neHEV-like particles (neHEV-LPs) have shown that neHEV-LPs adhere to cells via heparan sulfate proteoglycans (HSPGs) (17). Others have demonstrated that neHEV utilizes the transmembrane protein integrin α3 and eHEV the T cell immunoglobulin mucin domain 1 (TIM-1) receptor for host cell entry (18,19). Recently, the receptor tyrosine kinase EGFR (epidermal growth factor receptor) has been identified as a crucial co-factor for both neHEV and eHEV entry (20). Both HEV forms are internalized via a clathrin- and dynamin-2-dependent, receptor-mediated endocytosis (16,21,22). For eHEV, entry was suggested to involve trafficking to late endosomes and to be dependent on small GTPases Rab5 and Rab7, endosomal acidification and lysosome-mediated lipid degradation (16). Yet, the uncoating process, especially for neHEV, remains poorly understood.

Given the pivotal role of viral surface attachment to and entry into host cells in dictating viral host range, tissue tropism and pathogenesis (23), viral entry inhibitors offer a rational basis for the development of potent antivirals. In this study, the requirement for cellular proteases during neHEV and eHEV infection on viral entry were examined. Our results suggest that cysteine proteases of the cathepsin (CTS) family, in particular cathepsin L (CTSL), are critical host factors required during both neHEV and eHEV entry and provide a promising target to impede virus uptake into permissive cells.

## Methods

### 2.1 Compounds and reagents

Telaprevir (HY-10235) was purchased from MedChem Express or from Selleckchem (S1538). Leupeptin hemisulfate (HY-18234A), Pepstatin (HY-P0018), Aprotinin (HY-P0017), CA-074 (HY-103350), Brensocatib (HY-101056), Petesicatib (HY-109069) and E64d (HY-100229) were all purchased from MedChem Express. K11777 was purchased from both MedChem Express (HY-119293) or Biomol (AG-CR1-0158-M005), whereas CAA0225 (219502) was purchased form Sigma Aldrich. Compounds were dissolved in dimethyl sulfoxide (DMSO) or in a 50% DMSO/H_2_O mixture (Aprotinin) at a concentration of 25 mM, aliquoted and stored at -80 °C.

### 2.2 Eukaryotic cell culture

Hepatoma cell lines Huh7.5 and HepG2 were grown in Dulbecco’s modified Eagle’s medium (DMEM) (Invitrogen, Karlsruhe, Germany) supplemented with 10% fetal calf serum (FCS) (GE Healthcare), 100 µg/mL of streptomycin, 100 IU/mL of penicillin (Invitrogen), 2 mM L-glutamine and 1% nonessential amino acids (Invitrogen) (DMEM complete) at 37 °C in a 5% (v/v) CO_2_ incubator. Hepatoma cell line HepG2/C3A cells was grown in Minimum Essential Medium (MEM) (Invitrogen, Karlsruhe, Germany) supplemented with 10% ultra-low IgG FCS (GE Healthcare), 100 μg/mL of Gentamycin, 1% sodium pyruvate (Invitrogen), 2 mM L-glutamine (Invitrogen) and 1% nonessential amino acids (Invitrogen) (MEM complete). Commercially obtained human CTSL (Cathepsin L/MEP) knockout HEK-293T cells (abcam; ab266521) and human wild-type HEK293T cells (abcam; ab255449) were maintained in DMEM complete supplemented with 1 µg/mL puromycin or DMEM complete, respectively. HepaRG cell line was passaged and maintained in William’s E (Invitrogen) supplemented with 10% FCS (Invitrogen), 100 µg/mL Streptomycin (Invitrogen), 100 U/mL Penicillin (Invitrogen), 2 mM GlutaMax (Invitrogen), 5 µg/mL Insulin (Sigma-Aldrich, Taufkirchen, Germany) and 50 µM hydrocortisone hemisuccinate (Sigma-Aldrich) (HepaRG medium). For differentiation into mature hepatocytes, cells were seeded at a density of 50,000 cells/well of a 24-well plate and grown for at least two weeks until forming confluent layers. Medium was exchanged twice a week. Upon forming confluent layers, differentiation was triggered by adding 1.8% Hybri-Max DMSO (Sigma-Aldrich) to the HepaRG medium. Cells were passaged for at least two more weeks in the presence of 1.8% DMSO with two medium exchanges per week before performing experiments. Successful differentiation into cholangiocyte-like and hepatocyte-like cells was tested validated by immunostaining against albumin (Agilent, Cat. Nr. A0001). Primary human hepatocytes (PHHs) were purchased from Primacyt (Schwerin, Germany) as cryopreserved hepatocytes and thawed according to the manufacturer’s instructions (see Extended Table S1). PHHs were seeded on 24-well plates and kept in Human Hepatocyte Maintenance Medium (HHMM, Primacyt). Donors were serologically tested negative for following infectious diseases: HIV, hepatitis B and C, and SARS-CoV-2. Patient informed consent was obtained by Primacyt, as stated on their website.

### 2.3 Plasmids preparation, in vitro transcription and electroporation

Plasmids containing either the full-length viral genome of the Kernow-C1/p6 virus isolate (24,25) or the 83-2-27 isolate (26) (a kind gift from Takaji Wakita) were utilized for infectious viral particle production, and a plasmid encoding the sequence of the assembly-deficient subgenomic *Gaussia* luciferase reporter replicon (Kernow-C1/p6 strain with a truncated ORF2 replaced with a *Gaussia* luciferase gene) was used in replication assays. *In vitro* transcription and electroporation into cells and was performed as previously described (27,28). In brief, 5 µg *in vitro* transcripts were transfected into 5,000,000 cells resuspended in 400 µL cytomix containing 2 mM ATP and 5 mM glutathione using a Gene Pulser Electroporator (Bio-Rad, Munich, Germany). Cells were immediately transferred into 12 mL fresh medium and seeded in 96-well plates at density of 20,000 cells/well for replication assays and 12 mL/10cm dish for virus production.

### 2.4 Production of cell culture-derived HEV (HEV_CC_)

Infectious HEV_CC_ particle production was performed according to Todt *et al.* and Meister *et al.* (27,28). In brief, HepG2 cells were electroporated with *in vitro* transcripts from either the full-length Kernow-C1/p6 or the 83-2-27 strains. To obtain extracellular enveloped HEV_CC_ (eHEV_CC_), supernatant was collected seven days post transfection and stored at 4 °C until usage. Intracellular non-enveloped HEV_CC_ (neHEV_CC_) was harvested from cell lysate. This was achieved by trypsinizing cells, neutralizing them in DMEM, centrifugation at 200 × g for 5 minutes and resuspension in 1.6 mL per transfection. After three freeze and thaw cycles in liquid nitrogen and on ice, cells were centrifuged at 10,000 × g to separate the virus solution from cell debris. The cleared supernatant was aliquoted and stored at -80 °C.

### 2.5 HEV infection assays and compound titration

For HEV infection assays, cells were seeded at a density of 15,000 cells/well (HepG2, HepG2/C3A) and 5,000 or 8,000 cells/well (Huh7.5) of a 96-well plate and allowed to adhere overnight. Next day, medium was aspirated and exchanged with 50 µL medium that contained a dilution series of the compound of interest at 2x final concentration (TLV: 3-fold serial dilution starting at 10 µM. Aprotinin, Pepstatin, Leupeptin, K11777, CA-074, Brensocatib, Petesicatib, CAA0225 and E64d: 10-fold serial dilution starting at 10 µM). Each concentration was investigated in triplicates. HEV_CC_ was diluted in assay medium to reach a MOI of 0.1–2 in 50 µL total volume and added to the cells. Cells were incubated for 3–4 days (Huh7.5) or 4 days (HepG2; HepG2/C3A) before the cells were prepared for immunofluorescence staining and microscopy analysis. Differentiated HepaRG cells were infected with neHEV_CC_ and incubated in the presence of triplicate 10-fold serial dilutions of K11777 at concentrations ranging from 1 pM to 10 µM for 4 days. DMSO served as vehicle control. Primary human hepatocytes (PHH) were treated with 0.1, 1 and 10 µM K11777 for three days until immunofluorescence assay. Treatment with DMSO and 25 µM RBV served as a vehicle control and a positive control, respectively.

### 2.6 Replication assay

Hepatoma cells were transfected with Kernow-C1/p6 subgenomic replicon as described in 2.3 and incubated in the presence of 10 µM TLV, Aprotinin, Pepstatin and Leupeptin or 1 µM K11777. Supernatants were sampled 4, 24, 48 and 72 h post electroporation and stored at 4 °C until luminometer reading. Fifty micromolar of the replication inhibitor RBV served as a positive control.

### 2.7 Gaussia luciferase assay

*Gaussia* luciferase activity was measured by adding 20 µL of harvested cell culture supernatant per well on a 96-well LUMITRAC 600 plate, followed by the addition of Coelenterazine substrate and the detection of luminescence using a Centro XS3 LB 960 luminometer (Berthold Technologies, Bad Wildbad, Germany). The microplate reader was set to dispense 50 μL of substrate, followed by shaking for 2 s and reading for 5 s. Each condition was performed in triplicates.

### 2.8 Time-of-drug addition assays

HepG2/C3A cells were seeded at a density of 15,000 cells per well in a 96-well plate and were allowed to adhere overnight. The following day, K11777 was added at 1 hour prior to infection (-1 h), the time point of virus infection (0 h) or at 2, 4, 8, 12, 16, 24, 48 and 72LJh after infection at a concentration of 0.1 µM. Cells were infected with Kernow-C1/p6-FL virus (MOI of 0.1). At 96 h post infection, ORF2 positive cells were visualized by immunofluorescence staining and microscopic analysis. In parallel experiments, RBV (25 µM) was used as a reference compound for inhibition of viral replication.

### 2.9 Purified cathepsin assay

Cathepsin Inhibitor Assay Kits for cathepsin S (ab185437), cathepsin B (ab185438) and cathepsin L (ab197012) were all purchased from Abcam (Cambridge, UK) and performed according to the manufacturer’s instructions. In brief, purified enzymes were co-incubated in the presence of either different concentrations of compound of interest, a vehicle control, or a supplied known inhibitor as negative control. After incubation, substrate was added and production of fluorescent substrate was measured with a TECAN Infinite 200 Pro reader (Tecan Group Ltd, Männedorf, Switzerland) in kinetic mode measuring every 2 minutes at Excitation/Emission 400/505 nm. Reaction rate of vehicle control was normalized to 1, while reaction rate of the negative control was normalized to 0.

### 2.10 Immunofluorescence assay and microscopy

Cells were fixed in 3% paraformaldehyde at room temperature for at least 20 minutes and washed three times with PBS. Permeabilization was achieved using 0.1% Triton X-100 for 5 minutes, followed by three additional PBS washes. Cells were blocked in 5% horse serum in PBS for 1 hour at room temperature. HEV ORF2 capsid protein was stained with rabbit anti-HEV ORF2 polyclonal antibody #4086/#2101 (kind gift of Prof. Rainer G. Ulrich, diluted 1/5000 in blocking solution) overnight at 4 °C. Secondary antibodies were Alexa 488-labeled goat anti-rabbit IgG (Invitrogen A11008) or Alexa 488-labeled donkey anti-rabbit IgG (Invitrogen A21206), used at 1/1000 dilution in 5% horse serum-PBS, and incubated for 1–2 hours at room temperature in the dark. Nuclei were stained simultaneously with DAPI (1 µg/mL). Images were taken with a standard wide-field fluorescence microscope (Keyence BZ-X800E) with a 4 × 0.75-numerical aperture (NA) air-objective. DAPI (358 nm) and ORF2 (488 nm) signals were acquired sequentially by using the BZ-X Filter DAPI and BZ-X Filter GFP, respectively. Infections were quantified either by counting foci or, in the event of too many foci to count manually, by determining the percentage of positive cells using cell profiler (29).

### 2.11 Cell viability assays

Cell viability was determined by adding 0.5 mg/mL 3-(4,5-dimethylthiazol-2-yl)-2,5-diphenyltetrazolium bromide (MTT) (Sigma) substrate to cells and subsequent incubation at 37 °C and 5% CO_2_ for 1–2 h. Medium was removed and 50 μL of DMSO was added to each well. The absorbance of each well was read on a microplate absorbance reader (Tecan Group Ltd, Männedorf, Switzerland) at 570 nm. Cells treated with 70% ethanol for 10 min served as background control. Cell viability of PHH was determined using the CytoTox 96^®^ Non-Radioactive Cytotoxicity Assay (Promega). Lactate dehydrogenase release in the supernatant was determined 72 h post infection by transferring 50 µL supernatant to a 96-well plate, followed by the addition of an equal volume of CytoTox 96^®^ Reagent to each well. The reaction was incubated for 30 minutes and stopped by the addition of Stop Solution. Absorbance signal was measured at 492 nm in a plate reader (Tecan). Each condition was performed in triplicates.

### 2.12 Cathepsin expression

Normalized mRNA expression of Cathepsin family members in both PHH and Huh-7.5 cells was extracted from RNA-seq data reported in previous studies (30–32). For HepG2 cells, total RNA was extracted from four individual wells of seeded cells. The contents of each well were processed individually using a Machery Nagel kit according to the manufacturers protocol, and PolyA+ mRNAs were sequenced using the Illumina NextSeq550 platform. Normalized RNA-seq quantification of Cathepsin mRNA expression in HepG2 cells was determined after mapping to the Hg38 genome scaffold. Publicly available single cell RNAseq data was aquired using the R package CuratedAtlasQueryR (https://github.com/stemangiola/CuratedAtlasQueryR). The data was filtered for adult, non-diseased liver tissue which was aquired using the 10x genomics platform. Expression counts were normalized as counts per million.

### 2.13 Statistical analysis

Dose-dependent inhibition of infection was plotted and adjusted to a non-linear regression model using GraphPad Prism v9.3.1 for Windows (La Jolla, California, USA, www.graphpad.com). EC_50_, and CC_50_ were calculated using the four-parameter log-logistic model and statistical analysis was performed in GraphPad Prism v9.3.1 for Windows (La Jolla, California, USA, www.graphpad.com). Synergy was calculated with SynergyFinder (33).

## Results

### The HCV NS3/4A protease inhibitor telaprevir suppresses HEV infection in human liver cells

During co-infection studies with hepatitis C virus (HCV) and HEV (34), we noticed that the HCV NS3/4A serine protease inhibitor telaprevir (TLV), not only blocked HCV replication, but also inhibited HEV infection. To further investigate this unexpected effect, we evaluated the anti-HEV activity of TLV against HEV infection in a recently established HEV cell culture model (27) (**Fig. 1**). Human hepatoma cells were infected with non-enveloped cell culture-derived HEV (neHEV_CC_) in the presence of either TLV or a vehicle control (DMSO). TLV effectively inhibited neHEV_CC_ in Huh7.5 cells (**Fig. 1A, B**) in a concentration-dependent manner with a half-maximum effective concentration (EC_50_) of 1.5 µM (**Fig. 1C**). Notably, cell viability (**Fig. 1A, C**) was only slightly reduced. To specifically investigate a potential impact of TLV on HEV replication, we transfected subgenomic HEV-3 reporter replicon (Kernow-C1/p6-Gluc) into hepatoma cells and treated cells with 10 µM TLV. We found that HEV replication was not affected by TLV treatment (**Fig. 1D**), suggesting that TLV, though originally developed as an HCV protease inhibitor, influences the establishment of HEV infection rather than HEV RNA replication.

**Figure 1:**
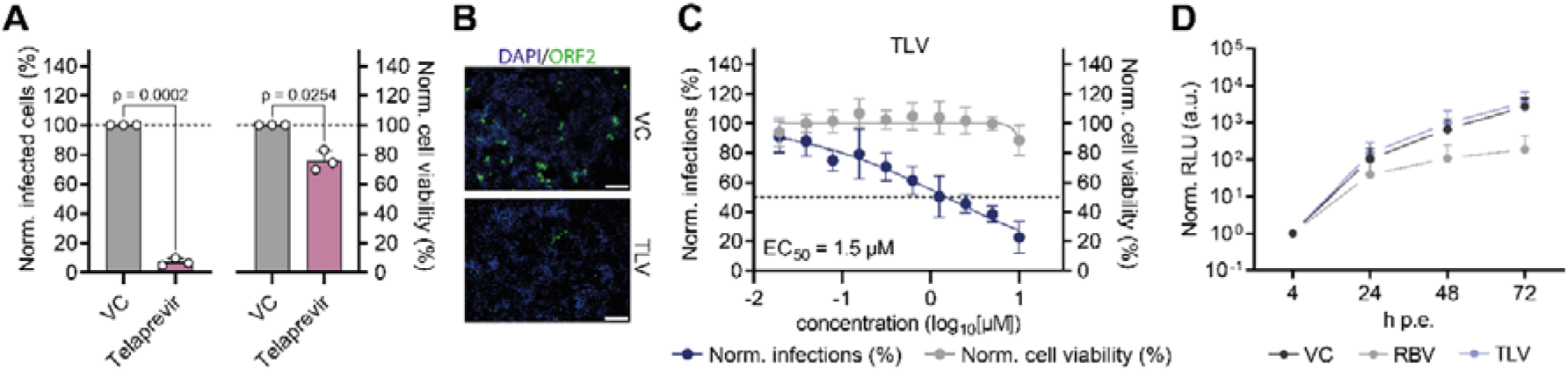
Role of HCV NS3/4A serine protease inhibitor telaprevir (TLV). (A) Huh7.5 hepatoma cells were inoculated with HEV in the presence of either 10 µM TLV or DMSO vehicle control (VC) for 4 days before fixation, immunofluorescence staining and analysis. Statistical significance was determined by two-tailed paired t-test. p-values >0.05 were considered to be not significant. (B) Representative immunofluorescence images illustrate the impact of telaprevir on the infectivity of non-enveloped cell culture-derived hepatitis E virus (neHEV_CC_). ORF2 = green; DAPI = blue; scale bar = 200 µm. (C) Dose-dependent neHEV_CC_ inhibition by TLV. Normalized infections in percent (%) were fitted by four-parameter log-logistic model to determine half-maximum effective concentration (EC_50_). (D) Replication capacity of Kernow-C1/p6 strain subgenomic replicon during treatment with TLV. Depicted are normalized relative light units (RLU) measured 4, 24, 48 and 72 hours post electroporation (h p.e.). Ribavirin (RBV) and dimethyl sulfoxide (DMSO) were employed as positive and negative controls, respectively. The depicted values represent means +SD (A, D) or ± SD (C) from three independent experiments.

### Leupeptin, a cysteine and serine protease inhibitor, suppresses HEV infection

Given that TLV is an HCV protease inhibitor, but HEV lacks structural proteins with known protease activities, we hypothesized that HCV protease inhibitors might target cellular-encoded proteases essential for HEV infection. To test this, we evaluated the effects of different broad-spectrum protease inhibitors (aprotinin, pepstatin and leupeptin) on HEV infection through dose-response assays (**Fig. 2A).** Notably, neither aprotinin, a serine protease inhibitor (EC_50_ > 10 µM), nor pepstatin, an aspartic protease inhibitor (EC_50_ > 10 µM), reduced HEV infectivity (**Fig. 2B, C**). In contrast, the cysteine and serine protease inhibitor leupeptin, displayed a concentration-dependent inhibitory effect on HEV infectivity (EC_50_ = 1.7 µM), implying a crucial role of cysteine proteases during HEV infection (**Fig. 2D**). Importantly, all tested protease inhibitors were non-cytotoxic (EC_50_ > 10 µM for aprotinin, pepstatin and leupeptin) (**Fig. 2E**). To specifically investigate a potential impact of the different protease inhibitors on HEV replication, we transfected hepatoma cells with subgenomic HEV-3 reporter replicon (Kernow-C1/p6-Gluc) and treated with 10 µM of protease inhibitors. None of the protease inhibitors reduced HEV replication (**Fig. 2F**), further indicating that cellular cysteine proteases are involved in the establishment of HEV infection, but not HEV RNA replication.

**Figure 2:**
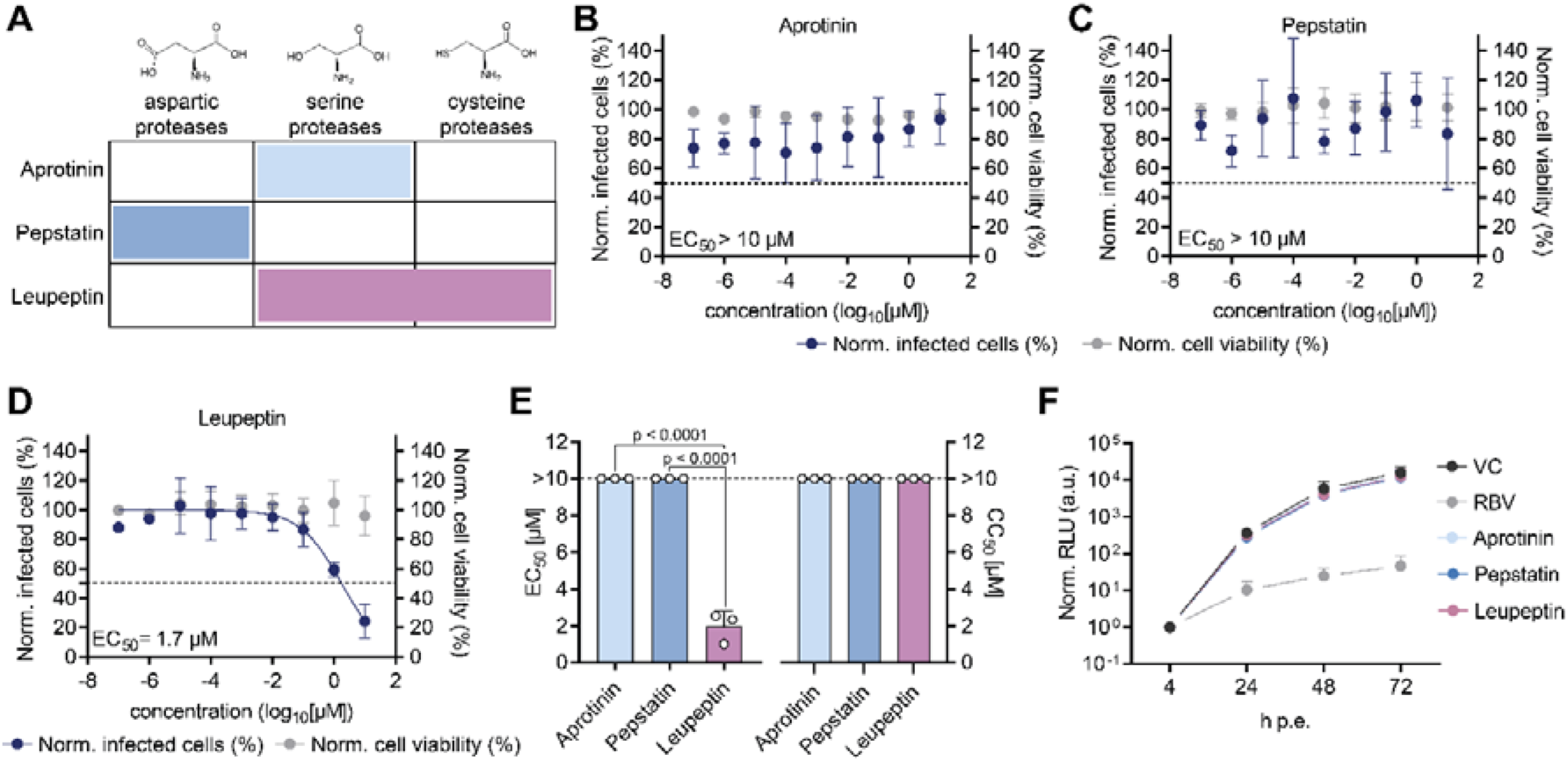
Role of cellular proteases in HEV infection. (A) Respective specificity of various broad-spectrum protease inhibitors. B-D) Hepatoma HepG2/C3A cells were infected with neHEV_CC_ and incubated with protease inhibitors or vehicle control (VC) for 4 days until immunofluorescence staining was performed against the HEV ORF2 capsid protein. Dose-dependent effects of aprotinin (B), pepstatin (C), and leupeptin (D) on neHEV_CC_ infections were plotted and fitted by four-parameter log-logistic model. (E) EC_50_ and CC_50_ values of aprotinin, pepstatin, and leupeptin in HepG2/C3A cells against infectious HEV Kernow-C1 p6 virus, determined by dose-response curves. To test the significance of mean differences, one-way ANOVA, followed by Dunnett multiple comparison test, was used. p-values >0.05 were considered to be not significant. (F) Impact of protease inhibitors on the replication of HEV subgenomic reporter replicon based on the Kernow-C1/p6 strain. Depicted are normalized relative light units (RLU) measured 4, 24, 48 and 72 hours post electroporation (h p.e.). Ribavirin (RBV) and dimethyl sulfoxide (DMSO) were employed as positive and negative controls, respectively. The depicted values represent means +SD (E, F) or ± SD (B-D) from three independent experiments.

### Cathepsin inhibitor K11777 impedes HEV infection

Cysteine proteases play pivotal roles in the entry process of several viruses, most notably Ebola (35) and SARS-CoV-2 (36). Interestingly, these cysteine proteases all belong to the CTS family, which are primarily localized in endo-lysosomal compartments and operates at acidic pH to digest incoming proteins. Blocking lysosomal acidification and thus CTS activity by ammonium chloride indeed strongly inhibited neHEV_CC_ infections (**Fig. 3A, B**). To ascertain if CTSs are crucial host factors for HEV infection, we first confirmed the inhibitory potency of the broad-spectrum CTS inhibitor K11777 against recombinant CTSs by a cell-free enzyme assay. The inhibitor displayed an EC_50_ of 0.9 µM, 3.3 µM and 92 nM for CTSS, B and L, respectively, with CTSL showing the highest sensitivity (**Fig. 3C**). In contrast, TLV required higher concentrations than K11777 to block CTS activity (**Fig. S1**). We next assessed the antiviral activity of K11777 in HEV infection experiments. In neHEV_CC_ infected cells, K11777 exhibited an EC_50_ of 232 pM and a CC_50_ > 10 µM (**Fig. 3D, G**), translating to a selectivity index of more than 10.000. Infection assays with the eHEV_CC_ isoform, showed a similar K11777 sensitivity compared to neHEV_CC_, with an EC_50_ of 18 pM for eHEV_CC_ (**Fig. 3E, G**), implying a similar dependency of both HEV isoforms on CTSs. Furthermore, infection with the wild-boar strain 83-2-27 demonstrated potent inhibition by K11777 with an EC_50_ of 550 pM (**Fig. 3F, G**), suggesting strain-independent antiviral activity of K11777. Also, an additive antiviral effect was observed in combination studies with RBV and K11777 (**Fig. S2**). Next, we investigated which viral life cycle step is perturbed by K11777. First, we determined replication capacity by HEV subgenomic reporter and found that similar to previously tested protease inhibitors, K11777 also did not affect virus replication (**Fig. S3**). Next, to elucidate the involvement of CTSs in the viral entry process, we performed time-of-drug addition assays in which K11777 was introduced at various stages: pre-infection (-1h), simultaneously during infection (0 h), and post-infection (2, 4, 8, 12, 16, 24, 48, 72h) (**Fig. 3H**). Notably, when introduced to cultures prior to infection as well as to cells already infected for up to 4 hours post-infection, K117777 retained its antiviral efficacy. However, a time-dependent loss in antiviral activity was observed when K11777 was administered for or beyond 8 hours post-infection, suggesting that K11777 targets the entry phase of the viral replication cycle and that CTSs are involved in virus entry. In contrast, a different pattern of viral inhibition was observed with the nucleoside analogue RBV, a well-known inhibitor of HEV RNA replication. The suppression of HEV was reduced only when RBV was administered following the initiation of viral RNA replication observed when RBV was added after onset of RNA virus replication (∼24 - 72 h p.i.), highlighting the distinct modes-of-action of K11777 and RBV in the HEV lifecycle.

**Figure 3:**
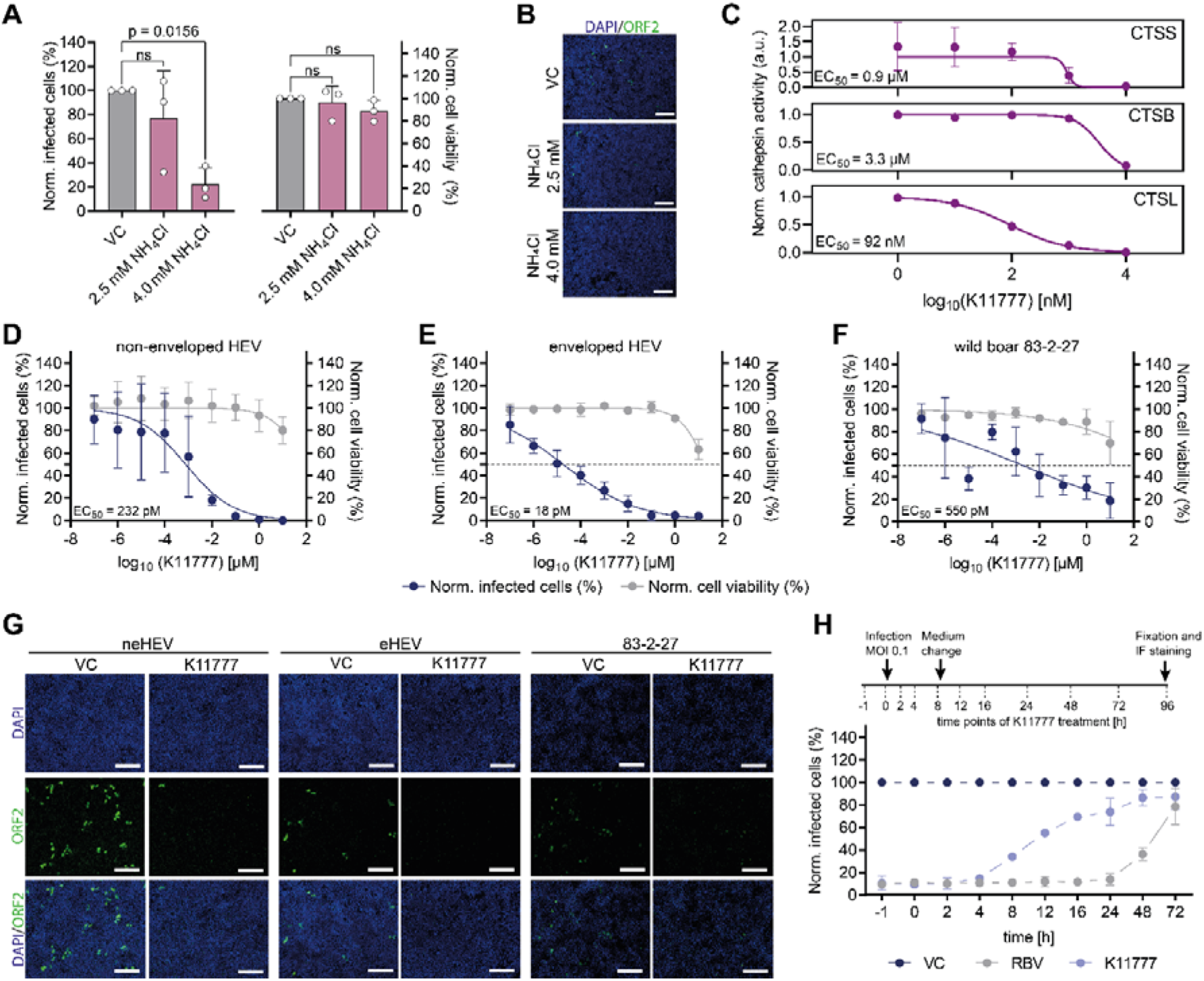
Cathepsin inhibitor K11777 impedes HEV late entry with nanomolar efficacy. (A) Blocking lysosomal acidification by NH_4_Cl inhibits HEV infection in hepatoma HepG2/C3A cells. Plotted are normalized (Norm.) infected cells in percent (%) and normalized cell viability in percent (%), determined for the treatment of 2.5 mM and 4 mM NH_4_Cl. DMSO served as vehicle control (VC). To test the significance of mean differences, one-way ANOVA, followed by Dunnett multiple comparison test, was used. p-values >0.05 were considered to be not significant (ns). (B) Representative immunofluorescence images of Kernow-C1/p6 virus infected and NH_4_Cl or DMSO (VC) treated cells. ORF2 = green; DAPI = blue.; scale bar = 200 µm. (C) Recombinant cathepsin S (CTSS), cathepsin B (CTSB) and cathepsin L (CTSL) were incubated with 10 µM K11777 in a cell free enzyme assay. D-F) Dose-dependency of K11777 on neHEV_CC_ infections (blue data points) and cytotoxicity (grey data points) in hepatoma HepG2/C3A cells. Cells were infected with HEV in the presence of K11777 or vehicle control for 4 days before fixation, immunofluorescence staining and analysis. Inhibition and cytotoxicity of Kernow-C1 p6 neHEVcc (D), eHEVcc (E) and wild boar 83-2-27 (F) strain by K11777. (G) Representative immunofluorescence images of K11777 treatment in HEV infected cells. ORF2 = green; DAPI = blue.; scale bar = 200 µm. (H, top) Experimental setup of the time-of-drug-addition assay. Hepatoma HepG2/C3A cells were inoculated with neHEV_CC_ at 0 h and treated with DMSO, K11777 or ribavirin (RBV) for 1 h (-1) prior to, during (0) and 2, 4, 8, 12, 16, 24, 48 and 72 hours after infection until fixation at 96 hours post infection. At 8 h after infection, inoculum was removed, and replenished with fresh medium containing drugs after several washes with PBS. (H, bottom) The inhibitory effect of 0.1 µM K11777 on HEV_CC_ infection when added at different time points pre- or post-infection is depicted by the blue curve. The broad-spectrum RNA virus inhibitor ribavirin (RBV; 25 µM) served as positive control (grey curve). Data presented represents meanLJ±LJSD from at two (C) or three independent experiments.

### Cathepsin L is required for HEV infection

Next, we aimed to identify which specific CTS(s) are involved in HEV entry. First, we assessed CTS expression in liver tissue using single-cell RNA-sequencing data from the human liver cell atlas, collected from nine healthy human donors (**Fig. 4A**). Notably, while most CTSs were expressed in the liver, CTSW, V, G and E had reduced expression. Kupffer cells exhibited the highest average expression, followed by hepatocytes and stems cells. Further examination of endogenous expression in primary human hepatocytes revealed prominent expression of CTSB, L, S and Z (**Fig. 4B**). A comparative analysis between HepG2 and Huh7.5 cells demonstrated only minor differences in RPKM levels and high levels of CTSB, C and L in both cell lines (**Fig. 4C**).

**Figure 4:**
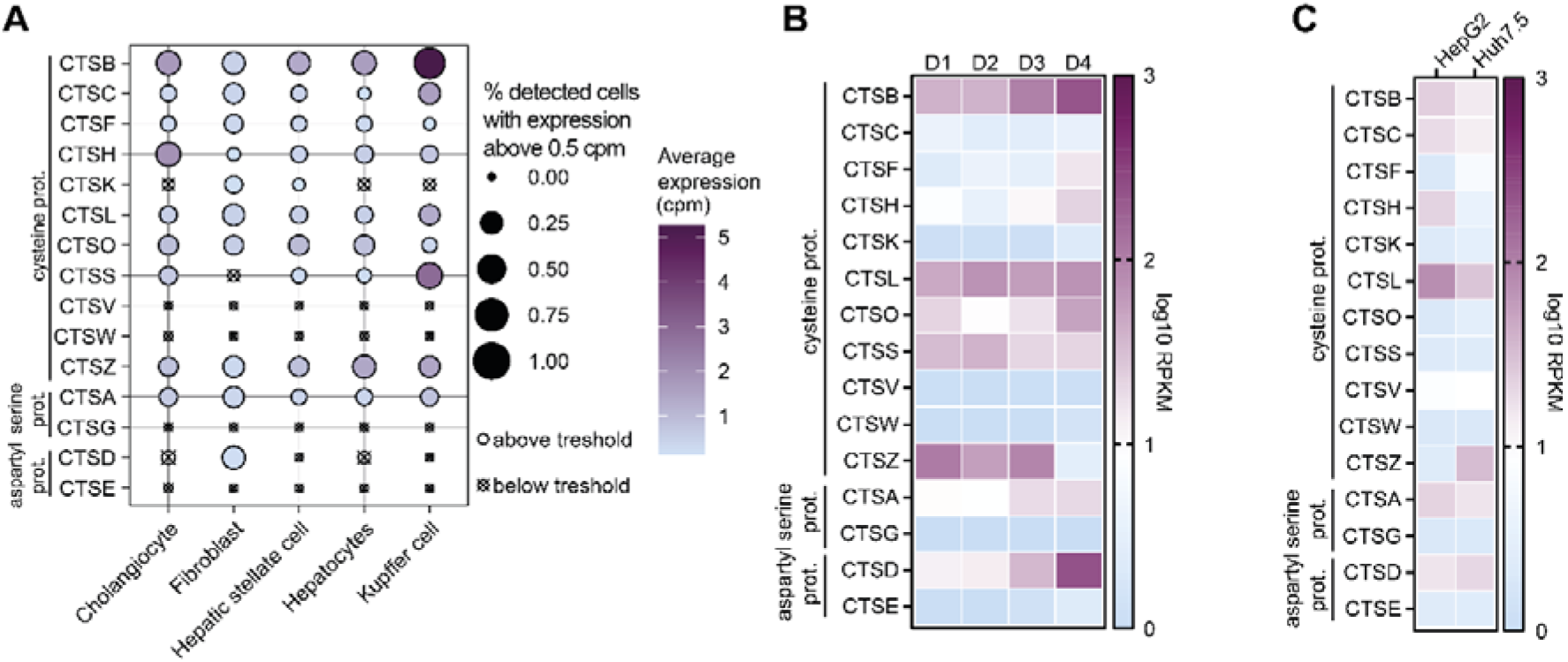
Cathepsins are endogenously expressed in primary human liver cells and standard hepatoma cell-lines. (A) Single cell analysis of liver-derived tissue. Expression data from single cell experiments was accessed via CuratedAtlasQueryR. The dotplot color visualizes average expression as counts per million (cpm) for different liver cell types. Dot size represents the proportion of cells that express a particular gene above 0.5 cpm. (B) Heat map of normalized transcript expression (reads per kilobase per million base pairs mapped [RPKM]) of CTSs in adult primary human hepatocytes and hepatoma cell lines HepG2 (mean of n=4) and Huh7.5 (mean of n=3).

To investigate the role of specific cysteine CTSs in HEV entry, HepG2/C3A cells were treated with specific cysteine protease CTS inhibitors (**Fig. 5A-E**). Notably, CTS inhibitors specifically targeting CTSB (CA-074) (**Fig. 5A**), CTSC (Brensocatib) (**Fig. 5B**) and CTSS (Petesicatib) (**Fig. 5C**) did not have a discernible impact on either cell viability or HEV infectivity. In contrast, two inhibitors, CAA0225 (**Fig. 5D**), a selective inhibitor of CTSL, and E64d (**Fig. 5E**), an inhibitor of both CTSL and CTSB, significantly inhibited HEV infection, characterized by an EC_50_ of 660 nM and 4 µM, respectively. To further confirm the importance of CTSL in HEV infection, commercially obtained human HEK293T CTSL homozygous knockout cells were infected with neHEV_CC_. Notably, CTSL knockout (CTSL KO) cells showed substantial reduction in HEV permissiveness, as evidenced by a reduced detection of the ORF2 capsid protein in immunofluorescence staining (**Fig. 5F, G**). In sum, these results demonstrate that CTSL is required for HEV infection.

**Figure 5:**
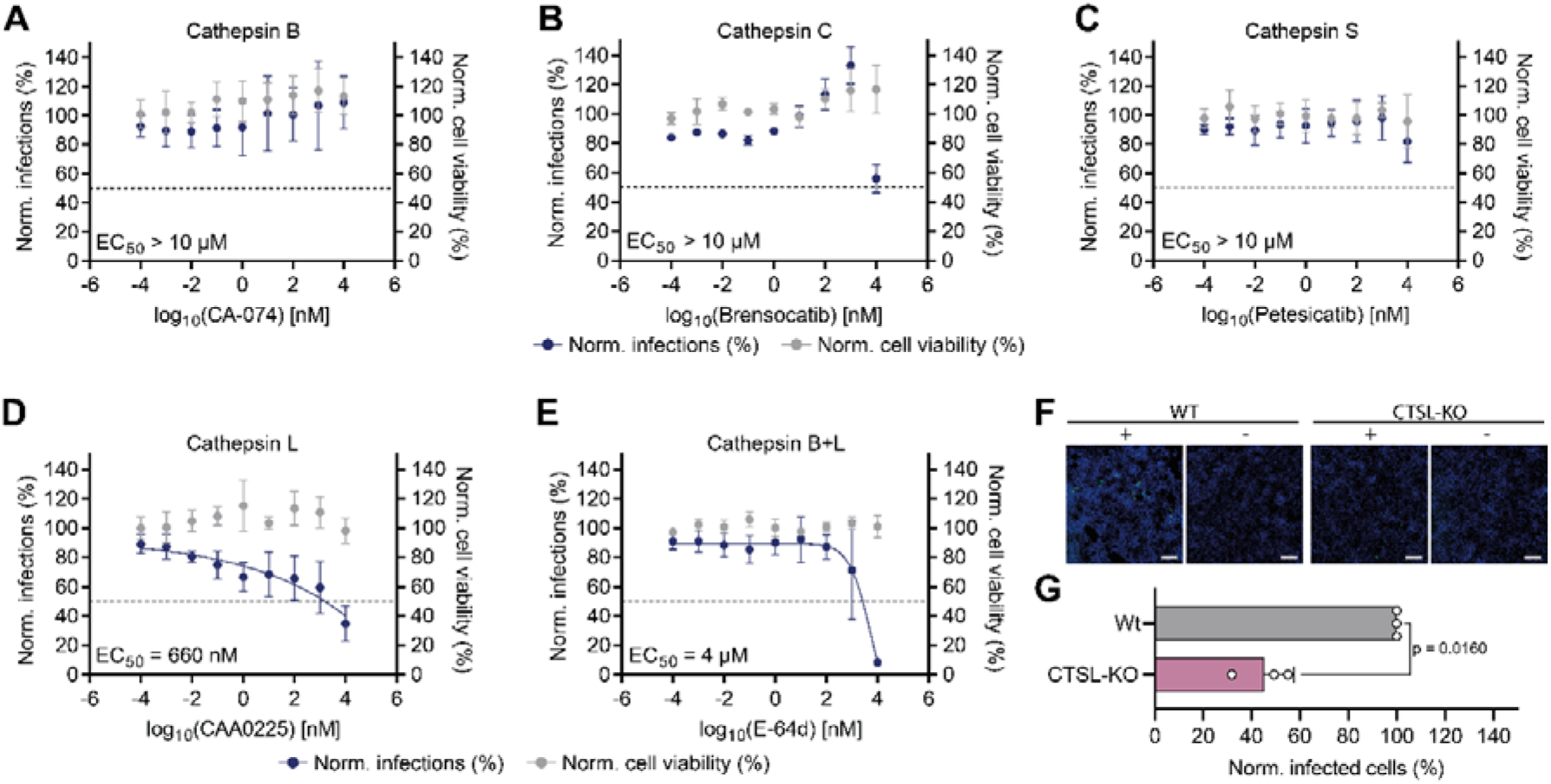
CTSL inhibitors block HEV infection *in vitro*. (A-E) Inhibition of HEV infection by different doses of CA-074 (A), Brensocatib (B), Petesicatib (C), CAA0225 (D) and E64d (E) and cell viability of cells treated with different doses of the drugs as indicated. (F, G) Effect of CTSL knockout on HEV infection, as determined by immunofluorescence staining against HEV ORF2 capsid protein. (F) Representative immunofluorescence images of HEK293T WT (abcam ab255449) and HE293T CTSL knockout cells (abcam ab266521). ORF2 = green; DAPI = blue; scale bar = 200 µm. (G) Data depicts means *±* SD of three independent experiments. To test the significance of mean differences, by two-tailed paired t-test was used. p-values >0.05 were considered to be not significant. Data presented represents meanLJ+LJSD from three independent experiments.

### Cathepsins are critical for HEV infection in HepaRG cells and primary human hepatocytes

To characterize the role of CTSs in a more authentic cell culture model, we assessed their significance for HEV infection in differentiated HepaRG cells (**Fig. 6A, B**). These hepatic cells emulate many features of mature hepatocytes such as high plasticity and bile canaliculi formation. We confirmed that K11777 inhibited HEV infections in HepaRGs (**Fig. 6A**) with an EC_50_ of 5.8 nM (**Fig. 6B**). To further address the importance of CTSs in primary liver cultures, we infected PHH with neHEV_CC_ in the presence of K11777. Our data revealed a dose-dependent HEV inhibition by K11777 (**Fig. 6C, D**) with almost no detectable ORF2 positive cells (**Fig. 6E**). Taken together, these data indicate that CTSs are critical for HEV infection in primary cells *ex vivo* and that HEV infection can be effectively restricted by the application of the K11777 inhibitor during inoculation.

**Figure 6:**
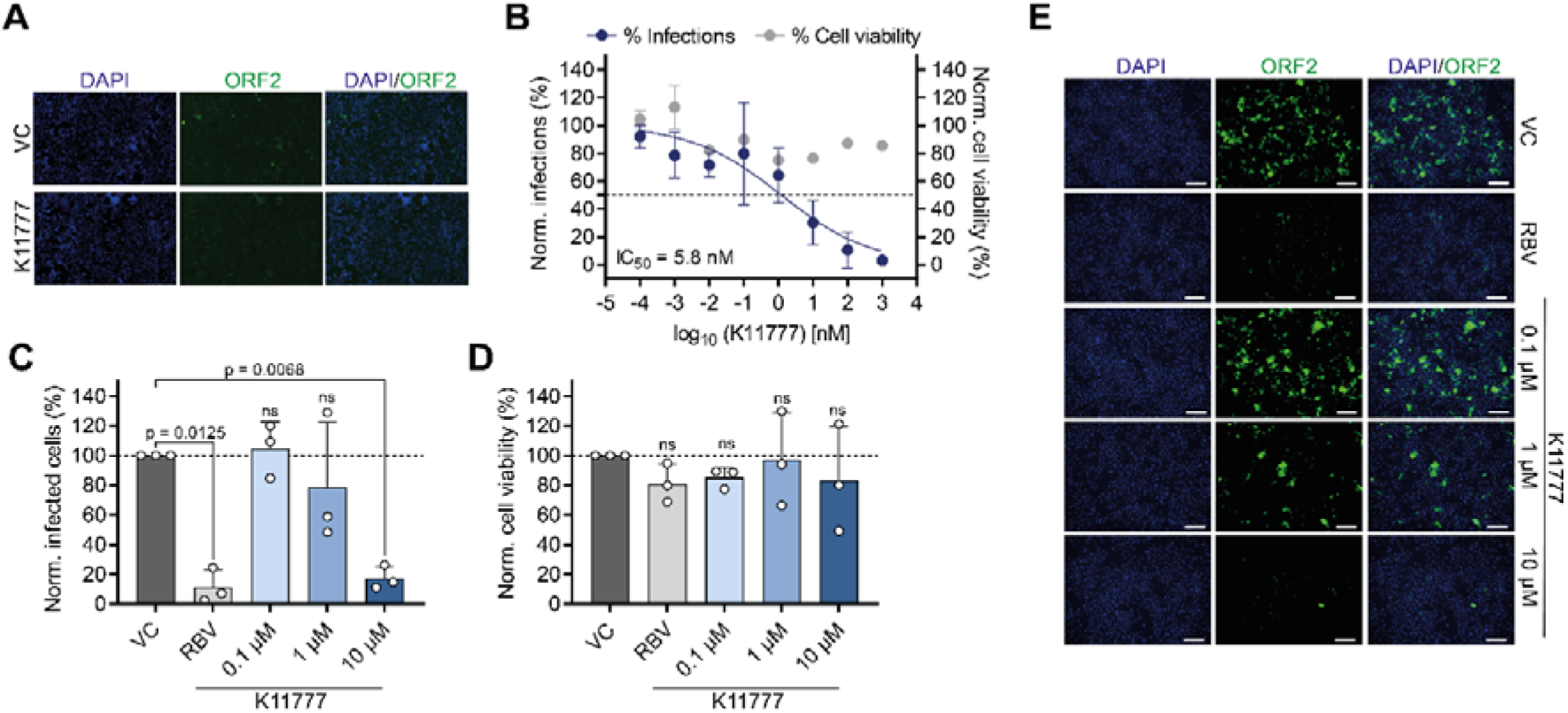
K11777 inhibits HEV infection in HepaRG cells and primary human hepatocytes. (A, B) Differentiated HepaRG cells were infected with neHEV_CC_ and incubated in the presence of K11777 or vehicle control (VC) for 4 days until immunofluorescence staining against ORF2 capsid protein. (A) Representative immunofluorescence images display the effect of 1 µM K11777 on infectivity of neHEV_CC_. ORF2 = green; DAPI = blue. (B) Dose-response curves of K11777 on neHEV_CC_ infections (blue datapoints) and cell viability (grey data points), as normalized to vehicle control, in HepaRG cells. Shown are means ± SD of three independent experiments. (C) Primary human hepatocytes were infected with Kernow-C1/p6 virus for 3 days and treated with 0.1, 1 and 10 µM K11777. Treatment with DMSO and 25 µM RBV served as negative and positive control, respectively. (D) Cell viability was determined by LDH release assay. Data depicts means + SD of three independent experiments. To test the significance of mean differences, one-way ANOVA, followed by Dunnett multiple comparison test, was used. p-values >0.05 were considered to be not significant (ns). (E) Representative immunofluorescence images. ORF2 = green; DAPI = blue; scale bar = 200 µm.

## Discussion

Although cell entry mechanisms used by viruses are highly diverse, many general principles of virus entry are conserved: viruses attach to target cells, internalize, cross a lipid bilayer, and deliver their nucleic acid to an intracellular site for replication. For HEV, these entry mechanisms remain poorly understood, largely due to the absence of a robust *in vitro* cell culture system to study HEV infections in the past decades. In this study, we utilized an advanced cell culture system to demonstrate the pivotal role of cysteine proteases, especially CTSs, in the entry of both neHEV_CC_ and eHEV_CC_. Moreover, we provided the first experimental evidence highlighting CTSL as an essential host factor for HEV entry, and demonstrate that the pan-cathepsin inhibitor K11777 significantly reduces HEV infection in both hepatoma and primary cells.

First, we observed that the HCV NS3/4A replication inhibitor TLV is effective against HEV infection, but not HEV RNA replication. This absence of observed reduction in HEV RNA replication upon TLV treatment led us to hypothesize that TLV does not target a putative HEV protease, but rather a host cellular protease. Indeed, current evidence indicates that in addition to direct viral targets such as HCV NS3/4A and SARS-CoV-2 M^pro^ (37,38), TLV might also affect indirect host targets, including CTSL (38). Another indication of CTS targeting by TLV stems from its structure. TLV is a linear α-ketoamide inhibitor that covalently binds to the catalytic serine (S139) of NS3/4A via it’s α-ketoamide warhead (39). These α-ketoamide motifs are frequently incorporated in lead molecules targeting serine and cysteine proteases (40). In fact, small molecules bearing an α-ketoamide warhead have been shown to inhibit CTSS potently at low nanomolar concentrations (41). In a similar vein, TLV’s α-ketoamide motif could potentially bind to CTSs, inhibiting their activity, however this warrants further investigation.

Next, we evaluated inhibitors targeting three primary classes of cellular proteases: cysteine, serine, and aspartyl proteases. Notably, we observed significant inhibition of HEV infection with leupeptin (EC_50_ = 1.6 µM), a combined serine and cysteine protease inhibitor. However, aprotinin and pepstatin, which are serine and aspartyl protease inhibitors respectively, did not demonstrate reduced infectivity. Similar to observations with telaprevir, HEV RNA replication was not impaired upon leupeptin treatment, further highlighting a role of cellular proteases in HEV infection rather than targeting any putative protease domains encoded by the virus genome. In addition, given that aprotinin and pepstatin inhibit trypsin and pepsin — two dominant proteases in the gastrointestinal tract — orally-transmitted viruses like HEV might have evolved to withstand the harsh gastrointestinal environment of the host, particularly the potential proteolysis of their structural proteins. In fact, it has been shown that the HEV capsid does contains trypsin digestion sites, however trypsin digestion is disrupted by the dimerization of ORF2, making it plausible that HEV remains uninhibited by these protease inhibitors (42). Interestingly, leupeptin has also been described as a potent “tight binding” inhibitor of CTSL (43), which aligns with our findings that highlight the significance of CTSL in HEV infection.

Our data also suggests, that neHEV_CC_ infection dependents on lysosomal acidification, a condition required for CTS activity. Treatment with NH_4_Cl, an agent that neutralizes endosomal pH, reduced HEV infectivity, which is in line with a recent study by Fu et al., who found that bafilomycin A, another endosomal acidification inhibitor was able to block neHEV_CC_ infection (44).

Based on the aforementioned results and the fact that CTSs are highly expressed and mainly lysosomal proteases involved in protein degradation (45), we investigated the role of different cysteine CTSs (CTSB, CTSC, CTSS, and CTSL). We found that the CTSL-specific inhibitor, CAA0225, reduced HEV infectivity with an EC_50_ of 660 nM, indicating a role of CTSL in HEV infection. Fittingly, we noted high CTSL expression in human hepatoma cells, primary human hepatocytes, and liver tissue, further highlighting its potential role in HEV entry.

To date, several viruses have been shown to rely on CTSs for effective cell entry, such as Ebola virus, SARS-CoV-2 and reovirus. For instance, CTSL and B, cleave the Ebola virus glycoprotein GP1, facilitating its interaction with cellular receptor(s) and viral entry (46,47), while the SARS-CoV-2 spike protein is cleaved by CTSB and L(48), initiating membrane fusion and uncoating of the viral RNA. CTSs crucial for reovirus entry proteolytically processes the capsid protein leading to structural rearrangements that eventually result in the release of transcriptionally active viral core into the cytoplasm (reviewed in (49). Reovirus can employ various CTSs (CTSB, CTSL, CTSS) for its processing (50,51), which is believed to contribute to its wide host tropism. By analogy, HEV might also use multiple different CTSs for its ORF2 capsid processing, explaining why HEV infection was not completely abrogated in CTSL knockout cells. Concurrently, this phenotype in CTSL knockout cells might be explained by the potential use of different entry strategies by HEV (similar to that observed for SARS-CoV-2). Also, HEV tropism is very broad, as evidenced by the fact that HEV is able to infect neuronal (52), placental (53) and human intestinal cells (54). Therefore, it will be of interest to determine if this broad cell tropism is in part a product of the ability to engage different CTSs.

Given that the data obtained in this study suggests that CTS activity plays a critical role in HEV entry into liver cells, targeting CTSs may represent an attractive targeting strategy for the development of new HEV antiviral drugs. The pan-specific CTS inhibitor, K11777, markedly reduced infectivity of both neHEV_CC_ and eHEV_CC_ as well as the wild boar strain 83-2-27 at pico- to nanomolar concentrations, confirming the importance of CTSs for both HEV isoforms and strains. Moreover, we have shown that K11777 inhibited a broad panel of CTSs and that this translated to a high potency in cell culture and most notably in *ex vivo* primary liver models. As we determined an additive antiviral effect when combined with RBV, whether combination therapy is able to increase efficacy and reduce resistance occurrence remains to be explored in future *in vivo* assays. K11777 has originally been developed against *Trypanosoma cruzei* (55), but has shown effective against several Filo- and Coronaviruses (56,57), including SARS-CoV-2 (58). Despite not being FDA-approved, it is orally available and currently undergoing a Phase-II clinical trial against SARS-CoV-2 under the name SLV213 (58), which is promising with respect to safety and tolerability. Future *in vivo* studies are essential to validate these encouraging findings and potentially expand the arsenal of effective HEV antivirals.

In conclusion, while HEV entry remains poorly understood, a process wherein HEV ORF2 undergoes CTSL-mediated proteolytic processing, resulting in viral RNA cytosolic release, appears plausible. However, further studies will be essential to define putative interactions between the HEV virion and CTSs. Finally, the CTS inhibitor K11777 emerged as a potent compound providing both excellent efficacy and selectivity *in vitro*.

## Acknowledgments

We kindly thank Prof. Rainer G. Ulrich, for the anti-HEV-ORF2 #4086 and #2101 antibody. We also thank Davide Durantel for kindly gifting the HepaRG cells.

## Author contributions

M.K., T.B., V.K., D.T. and E.S. designed research; M.K., T.B., J.J., J.A.S., performed research, M.K., T.B. and E.S. analyzed data; M.K., T.B. and E.S. wrote the original draft; M.K., T.B., J.J., J.A.S., A.G., R.J.P.B., V.L.D.T, V.K., Y.B., D.T. and E.S reviewed and edited the original draft.

## Conflicts of interest

The authors have no competing interests.

## Financial support and sponsorship

E.S. was supported by the German Research Council (STE 1954/12-1 and 1954/14-1), German Centre for Infection Research (DZIF, TTU 05.823_00) and by the Ruhr University Bochum InnovationsFoRUM (Project: Host Microbe Interactions, IF-018N-22). D.T. was supported by grants from the German Ministry of Education and Research (BMBF, project VirBio; 01KI2106). V.L.D.T was supported by the German Centre for Infection Research (TTU 05.823_00) and the German Research Council (DA 1640/3-1).

## List of Abbreviations

CTS: Cathepsin
Gluc: *Gaussia* luciferase
HCV: Hepatitis C virus
HEV: Hepatitis E virus
HEVcc: Cell culture derived HEV
KO: Knockout
PHH: Primary Human Hepatocytes
RBV: Ribavirin
RLU: Relative light units
WT: Wildtype

## Notes

### Competing Interest Statement

The authors have declared no competing interest.

