## Supplement Information for "Targeting cellular cathepsins inhibits hepatitis E virus infection"

*equal contribution

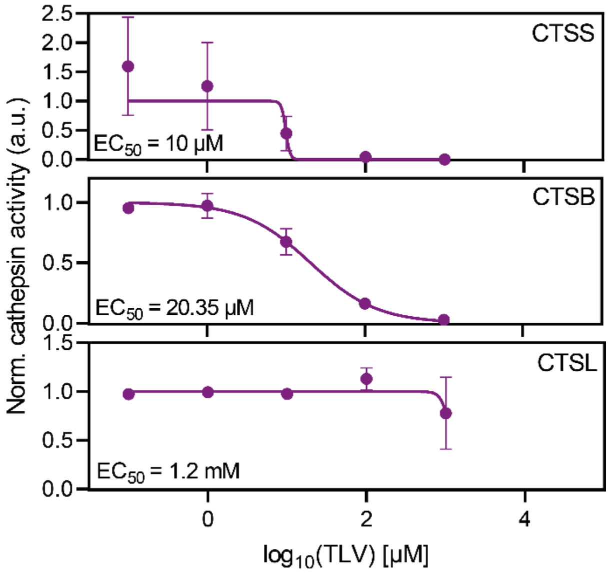


**Figure S1: TLV inhibits cellular cathepsins in cell free enzyme activity assay.** Recombinant cathepsin S, B and L were incubated with different concentrations of TLV and tested for their enzymatic activity in a cell free enzyme assay.

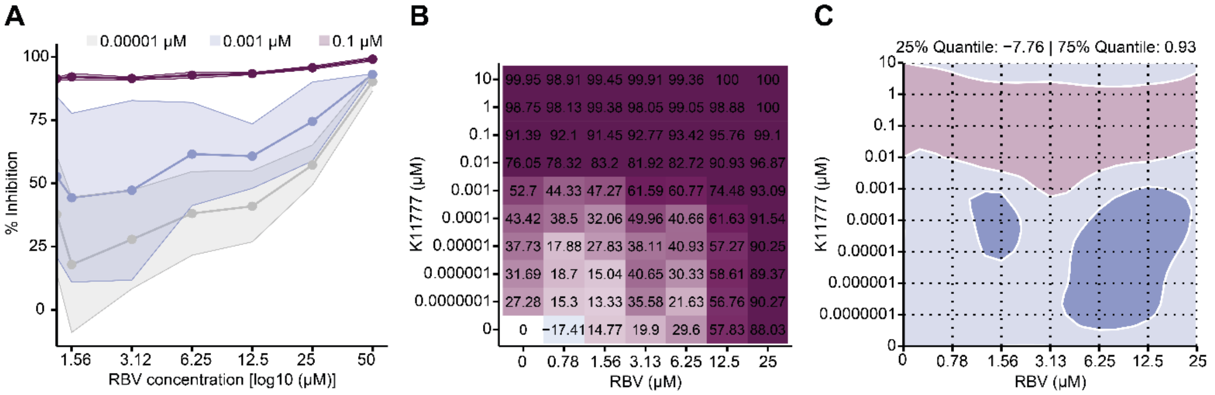


**Figure S2: *In vitro* effect of antiviral combinations on HEV infection in HepG2/C3A cells.** (A) HEV inhibition during treatment with 0.1 µM (red), 0.001 µM (blue) or 0.00001 µM (grey) K11777 and simultaneous titration of RBV (from 0 - 25 µM). (B) The dose-response matrix for RBV + K11777 are presented, depicted as the percent inhibition of viral infection in HepG2/C3A cells. (C) A two-dimensional map of synergy scores shown for the combination of K11777 with ribavirin (RBV). Synergy scores were based on ZIP synergy analysis determined with SynergyFinder. Determined ZIP score was -4.29. The ZIP score stands for the response beyond expectation in percentage. In the range of -10 < ZIP < 10, the compounds are likely to act in an additive manner, while a score ≥ 10 indicates synergism, and less than -10 shows antagonism. Data presented represent the means of three independent experiments.

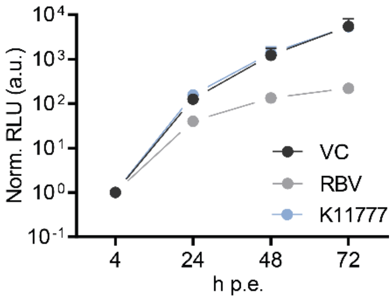


**Figure S3: Effect of 1 µM K11777 on replication of HEV subgenomic reporter replicon based on Kernow-C1/p6.** Depicted are normalized relative light units (RLU) measured after 4, 24, 48 and 72 hours post electroporation (h p.e.). Fifty µM Ribavirin (RBV) was used as a replication inhibition control. Dimethyl sulfoxide (DMSO) served as vehicle control.

**Extended Table S1: Donor Data of primary human hepatocytes used in this study.**

| Lot# | Species | Gender | Race | Age | BMI | Smoker | Alcohol use | Drug use | Pathology | Seeding density in 24 well |
|---|---|---|---|---|---|---|---|---|---|---|
| CHM2225-He-Z | human | male | caucasian | 73 | 27.99 | No | No | No | Hepatocellular Carcinoma | 210,000 cells/cm2 |
| NHM2354-HE-N | human | male | caucasian | 2 | / | No | No | No | Cardiac arrest | 233,000 cells/cm2 |
| BHum15052 | human | male | caucasian | 54 | 23 | No | No | No | Colorectal cancer hepatic metastases | 224,000 cell/cm2 |
